# Seasonal diets overwhelm host species in shaping the gut microbiota of Yak and Tibetan sheep

**DOI:** 10.1101/481374

**Authors:** Xiaojuan Wei, Fusheng Cheng, Hongmei Shi, Xuzheng Zhou, Bing Li, Ling Wang, Weiwei Wang, Jiyu Zhang

## Abstract

Host genetics and environmental factors can both shaping composition of gut microbiota, yet which factors are more important is still under debating. Yak (*Bos grunniens*) and Tibetan sheep (*Ovis aries*) are very different from the size and genetics. Nomadic Tibetan people keep them as main livestock and feeding them with same grazing systems, which provide a good opportunity to study the effects of diet and host species on gut microbiome. We collected fecal samples from yaks and Tibetan sheeps at different seasons when they were feed with different diets. Illumina data showed that major bacterial phyla of both animals are Bacteroidetes and Firmicutes, which agree with the previous reports. And the season effect had a higher impact on the gut microbiota than that of host species, though the animals are taxonomically distinguished each other at subfamily level. Since that the animal grazing differently at different seasons, this study indicated that diet can trump the host genetics even at higher taxonomic level. This finding provides a cautionary note for the researchers to link host genetics to the composition and function of the gut microbiota.

**Importance:** Yak and Tibetan sheep are very different from the size and genetics (from different sub-family). Nomadic Tibetan people keep them as main livestock and feeding them with same grazing systems, which provide a good opportunity to study the effects of diet and host species on the gut microbiota. Results indicated that diet can trump the host genetics even at higher taxonomic level. This finding provides a cautionary note for the researchers to link host genetics to the composition and function of the gut microbiota.

## Introduction

The high altitude makes the Qinghai-Tibetan Plateau become extreme harsh environments for the survival of mammalian species. There are two typical high-altitude ruminants, Yak (*Bos grunniens*) and Tibetan sheep (*Ovis aries*), being adaptively living in this harsh environments and turning to be nomadic Tibetan people’s livestock [1]. They are essential in providing food (milk and meat), transport (mainly yak), and fuel (feces of yak), shelter and clothes (skin and fur), and also fulfill various socio-cultural functions within the pastoral society.

As livestock, Yaks and Tibetan sheep are in the same grazing system or fed with the same feeding stuff, which provides a good opportunity to study the gut microbiota with different host species but similar diets. In addition, grazing systems in Qinghai-Tibet Plateau have seasonal changes in the different pastures with the different forage [2], especially between summer and winter. It’s a nice “treatment” that varied the diet to the Yaks and Tibetan sheep.

In the recent decade, intensive studies indicated that there are many factors can shape the composition of gut microbiota in mammals, including host genetics and diet [3-8]. Some reports showed that host genotype had a measurable impact on gut microbial community in both humans [9, 10] and mice [11]. But there are also reports showed that diet can overrule genotype differences in mouse gut microbiota [12], which mean that diet matters more than that of host genetics. We notice that the 5 inbred mouse strains in the experiments are belonging to the same species, and raise a question about how far phylogenetic distance of the host mammals can be masked by the diet.

In the current study, Yak and Tibetan sheep are belonging to the same family, namely Bovinae, but different subfamily, Bovinae and Caprinae, respectively (Wikipedia). We investigated gut microbial community at spring and autumn to test which factor has more impact in shaping the composition of gut microbiota, host species or diet.

## Results

### Variations of the gut prokaryotic community over the season and hosts

Illumina sequencing yielded a total of 4,363,232 raw reads of 16S rRNA gene sequences. After quality filtering, 3,021,303 valid sequences were clustered into 6,784 prokaryotic operational taxonomic units (OTUs) at 97% sequence identity level. Venn diagrams showed that the OTUs are mostly distinguished by season (Fig.1a). But to the functional genes, gut microbiota shared most of the genes regardless the seasons and the hosts (Fig.1b).

**Fig. 1.**
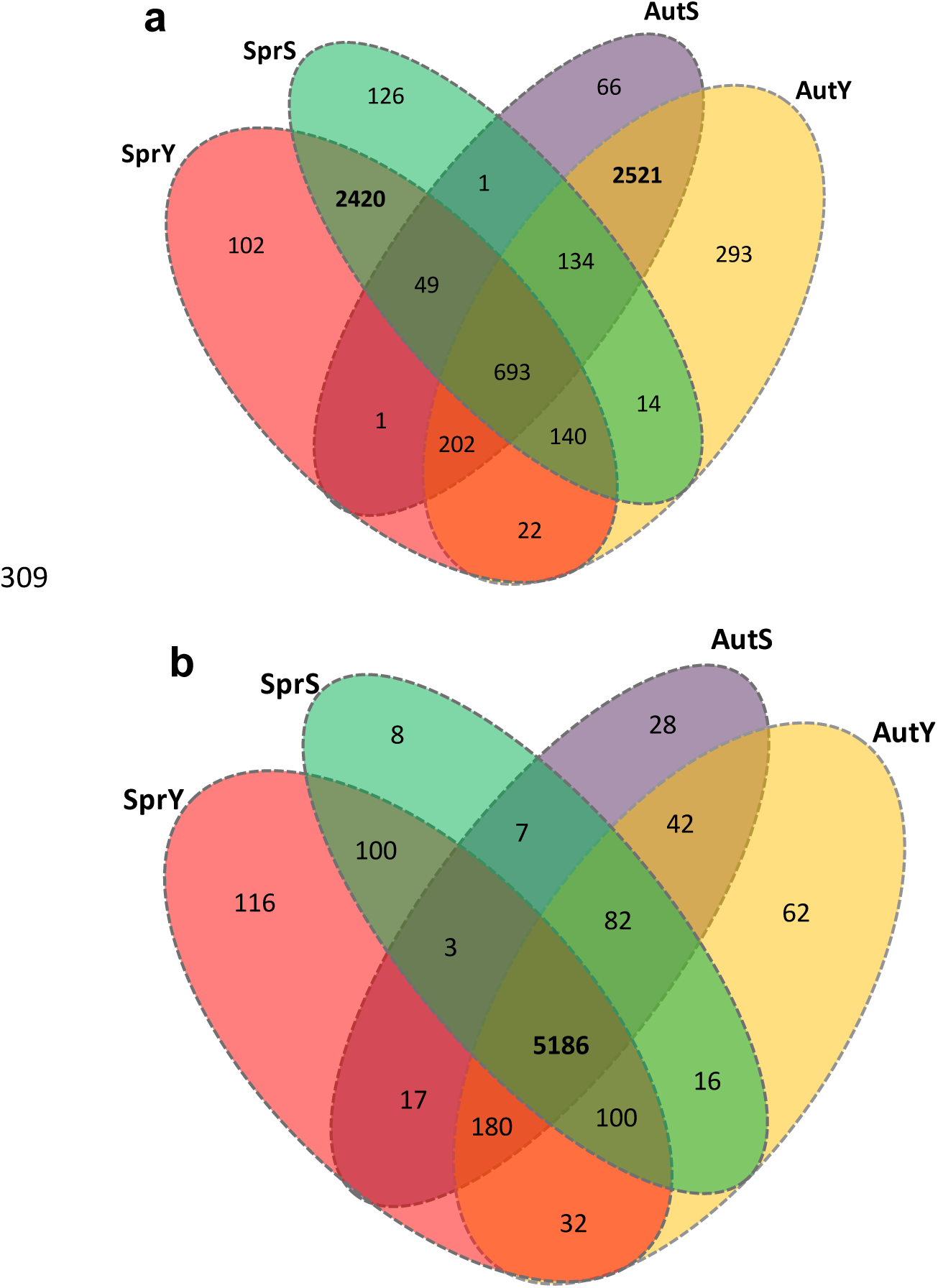
Venn diagrams of the taxanomic OTUs(a)and the KEGG functional genes(b)predicted by PICRUSt.

Overall, we identified 22 bacterial and 1 archaeal phyla in all investigated samples. Bacteroidetes is the most predominant phylum which averagely comprised 56% of the relative abundance. In spring, yaks and sheep had similar relative abundances of Bacteroidetes in their gut. In autumn, yaks had significantly higher abundance of Bacteroidetes than that of sheep (P < 0.001) (Fig. 2). But if group the samples with season only, there is no significant differences in the relative abundances of Bacteroidetes (P>0.1). Firmicutes is the secondly most predominant phylum, takes 38% of the total prokaryotic community in average. Firmicutes showed no significant variations when the samples grouped by either season or host, but the interactions between the seasons and hosts are significant, namely the changes in different seasons are different in the different hosts. Other phyla changed more by season than that by the host. At family levels, the variations were much stronger, especially by season (Fig.2). Bacteroidaceae and Rikenellaceae showed us in spring, while Prevotellaceae, BS11, and S24-7 proliferated during autumn. Fibrobacteraceae and Spirochaetaceae occur mainly in the gut of sheep and during the spring only.

**Fig. 2.**
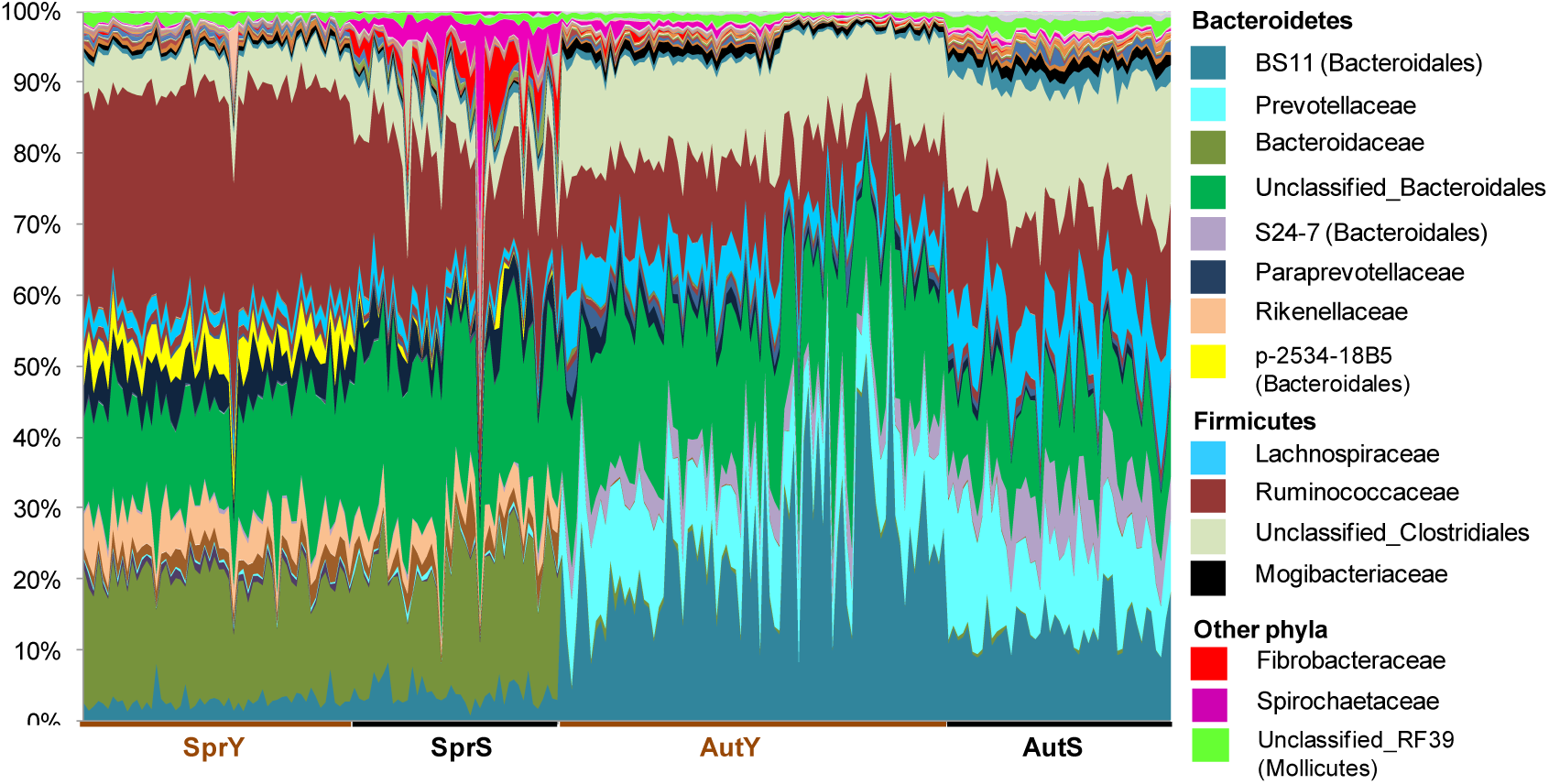
prokaryotic community composition at family level. The y-axis showed the values of the relative abundances of families. The x-axis is the samples which were grouped by host and season.

### Host and season effects on diversity indices

To the species richness (chao1), both host and season effects are highly significant. But there is no interaction between host and season effects: yaks always have more prokaryotic species in their gut than that of Tibetan sheep, and there are always more species in autumn than that of spring (Fig.3). Host effect is marginally significant to the evenness (Simpson) but season effect is highly significant. The interaction of the effects is also significant. To Shannon index, both host and season effects are not significant, but the interactions of the effects are significant (Fig. 3). Beta diversity is much higher between different seasons than that between different hosts; the later is barely higher than that within the same groups (Fig.4).

**Fig. 3.**
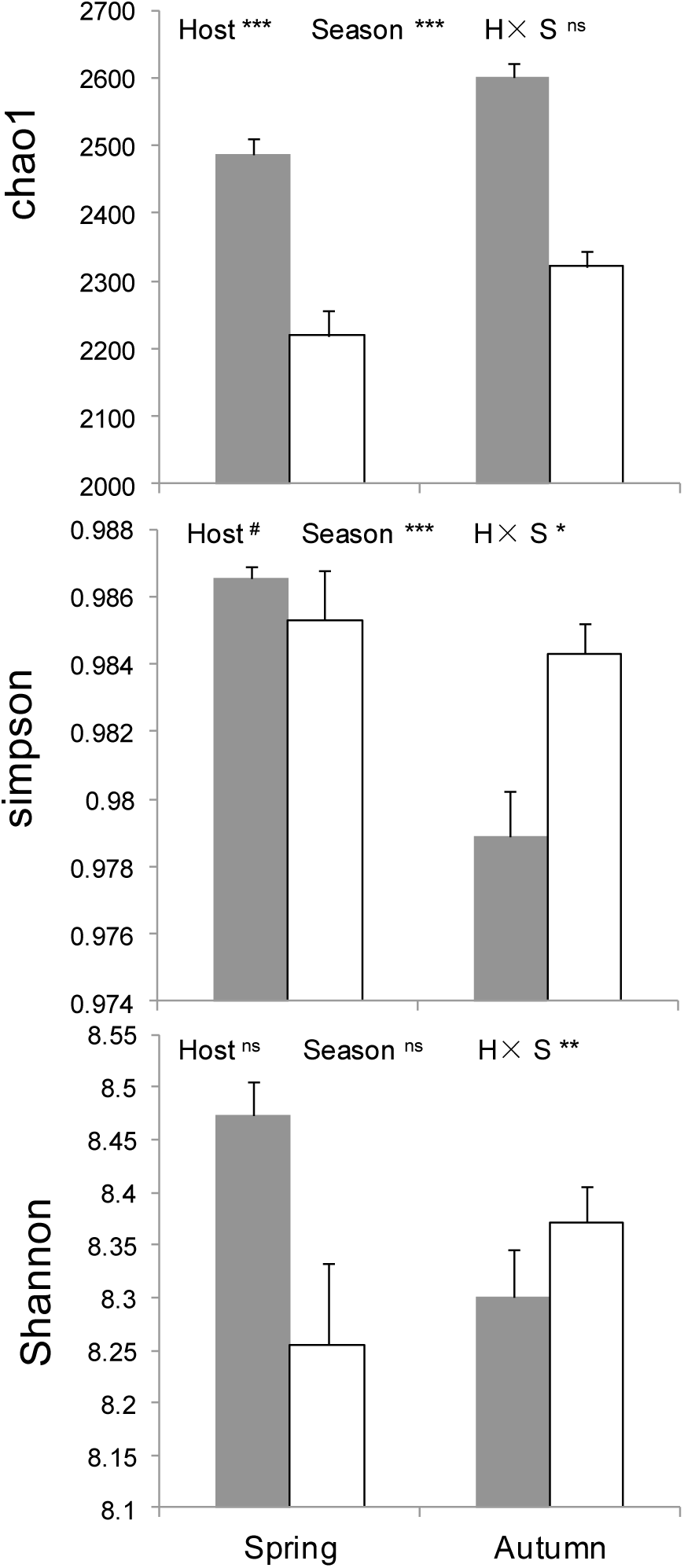
Two-way ANOVA analysis for alpha diversity indices (Mean ± SE) with host and season. Gray bars are data of yaks and white bars are Tibetan sheep. *, p<0.05; **, p<0.01; ***, p<0.001; #, p<0.1.

**Fig. 4.**
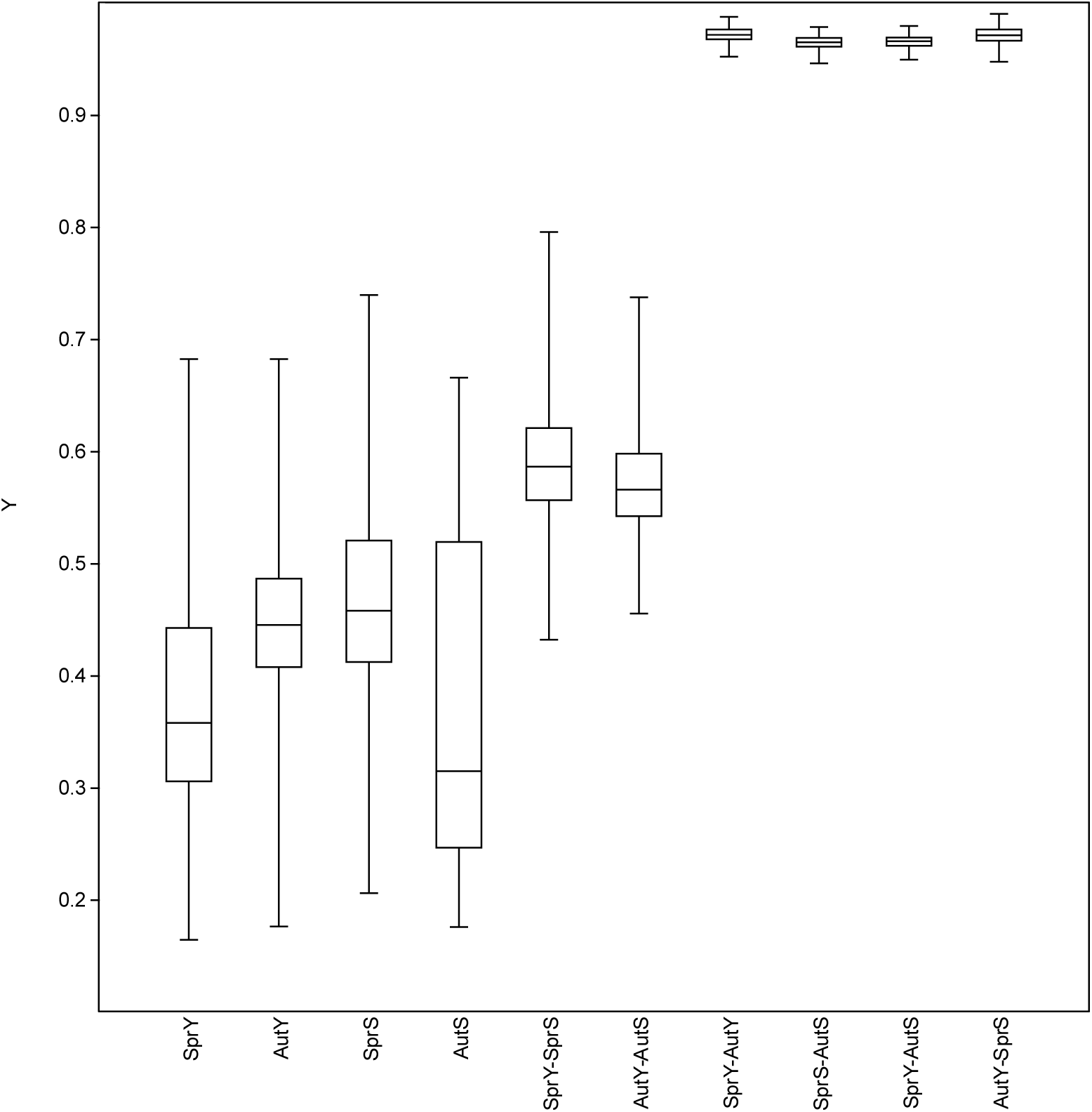
Beta diversity based Bray-cutis distances of the samples inside the groups and between the different groups.

### Species composition and functional genes composition

Principal co-ordinates analysis (PCoA) plots were generated to compare the composition of the microbial community among the hosts and seasons (Fig. 5). In PCoA plot of the prokaryotic community (Fig. 5a), PCo1 explained 57.4% of the variances and clearly divided the samples from spring and autumn, while PCo2 explained 7.1% of the variances and mainly differentiated the host species, which is effectively separated in autumn only. As for functional gene composition (Fig. 5b), PCo1 explained 70.6% of the variances and PCo2 explained 13% of the variances. But the groups cannot distinguish from each other clearly, though the results of permanova indicated the host effect, season effect, and the interaction between them are significant.

**Fig. 5.**
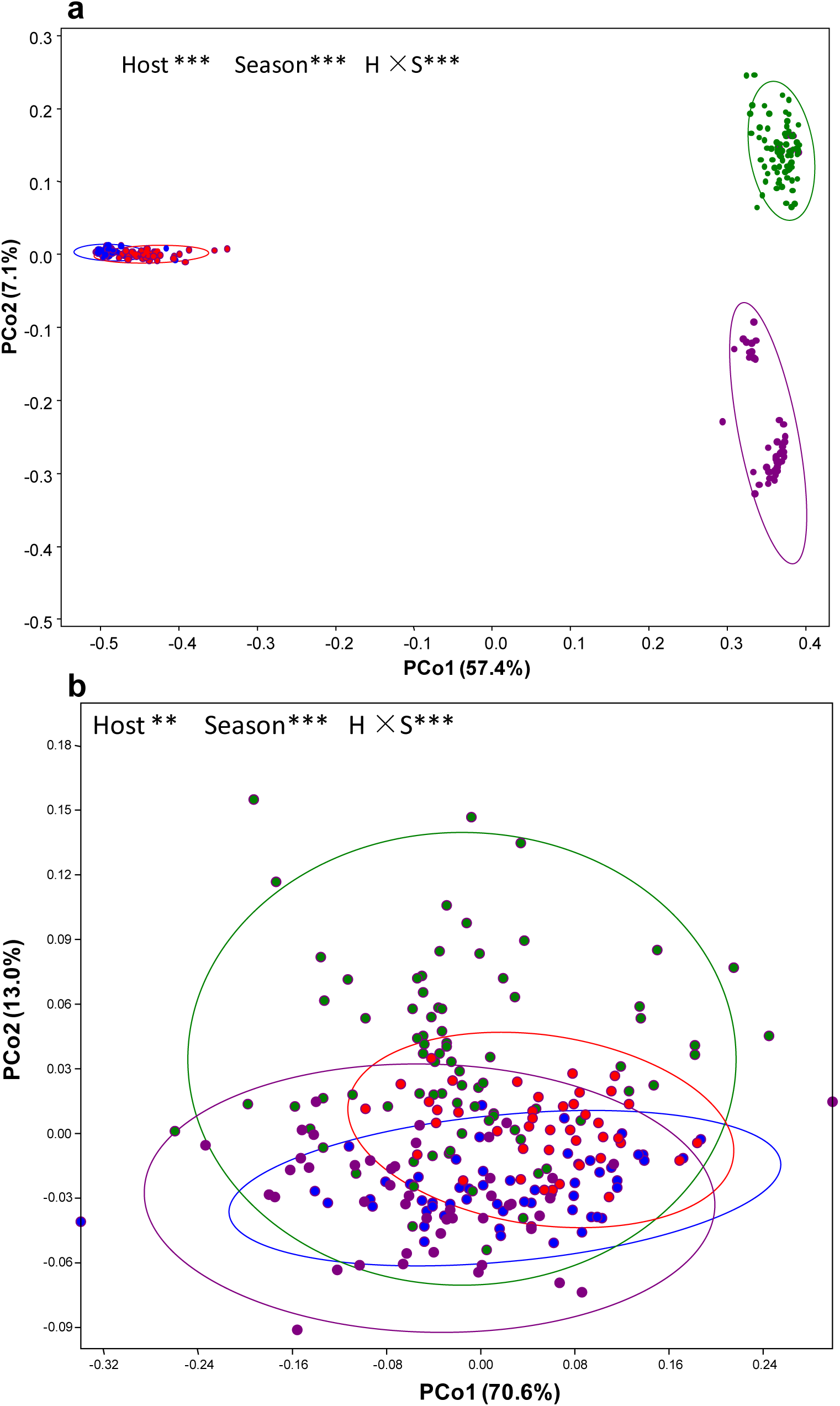
Principal coordinates analysis (PCoA) ordination of the taxanomic OTUs (a) and the KEGG functional genes (b) predicted by PICRUSt. Dots indicate one sample and the cycles are the 95% ellipses. Colors are as follows: blue = SprY; red = SprS; green = AutY; purple = AutS. Results of PERMANOVA are given in the higher right of each panel: **, p < 0.01; ***, p<0.001.

## Discussion

The gut microbiota of the mammals is acquired from the environment starting at birth. The assembly of the microbial community is largely shaped by environmental factors such as age, diet, lifestyle, hygiene, and disease state. Besides, the host genetics are also important to the composition of the gut microbiota. Subconsciously, researchers think that the host species will be more important than environmental factors in shaping gut microbiota, especially when the host species are very different taxonomically. Hence, it’s very rare to find the studies to directly compare the gut microbiota from different species of the animals.

Here, our results indicate that seasonal changes can overrule the variation come from host species, though yak and Tibetan sheep are very different taxonomically and also from the body size. There could be several explanations.

Firstly, both yak and Tibetan sheep are rumen animals. The rumen provides a strictly anaerobic environment where the microbes degrade plant fibers, nonfiber carbohydrates, and protein into volatile fatty acids and ammonia, which are used by rumen microbes as energy and nitrogen sources for their own growth. Therefore, rumen microbes could be possibly more similar than the gut microbes from elsewhere. In our study, gut microbiota in both animals is predominated by Bacteroidetes and Firmicutes, which were agreed by the previous reports of rumen microbiota of yaks [13, 14]. With another thought, the rumen microbiota could possibly be the starting of gut microbiota and did not change dramatically after coming out from the rumen. By the way, if it’s real, a deep understanding of microbial composition and variation is necessary to improve the welfare, health and production efficiency of ruminant livestock.

Secondly, in our study, the yaks and Tibetan sheep are always live together. Hence, the initial source of gut microbiota could come from the same environment. As is already known, early life events will be critical for gut microbiota development towards the adult microbiota. Lifestyle and diet will further influence the structure and function of gut intestinal microbiota. But in the studied animals, they have a very similar lifestyle and the same diet source. In our results, sheep and yaks had a nearly same composition of gut microbiota in the spring samples but distinguished from each other in autumn samples. The reason could be that, during summer and autumn, pasture grow more grass which allow the animal have diet selection, after all, sheep have different diet preference from that of yaks [2, 15-17]. However, in winter, there is no option but to eat the same food for survival.

Third, there could be a convergent evolution of gut microbiomes in yaks and Tibetan sheep due to the extremely harsh environment at the high altitude regions [1, 18]. When compared with their low-altitude relatives, cattle (*Bos taurus*) and ordinary sheep (*Ovis aries*), metagenomic analyses reveal significant enrichment in volatile fatty acids yielding pathways of rumen microbial genes in yaks and Tibetan sheep, whereas methanogenesis pathways show enrichment in the cattle metagenome. Analyses of RNA transcriptomes reveal significant upregulation in 36 genes associated with volatile fatty acids transport and absorption in the ruminal epithelium of yaks and Tibetan sheep. Which means, other than host genetics, long-term threaten of harsh environments will allow gut microbiome to be adaptive to help the host in health maintenance and survival. In other thought, though yaks and Tibetan sheep are very different in their own genetics, but inside their gut, microbiome could be more similar for adaptation of the high altitude.

Here, we also notice that the differences in the gut microbiota composition are mainly from the taxonomic aspect. By functional genes, both the host effect and season effect are not obvious. One possibility is that the variation of the composition is only some substitution of the microbes with the same functions. The other possibility is the prediction of PICRUSt might be partially inaccurate when applying to high altitude mammals, after all, the database developed for PICRUSt is lack of suitable data.

In summary, we find that diet can trump the host genetics even from different subfamily. This finding provides a cautionary note for ongoing efforts to link host genetics to the composition and function of the gut microbiota.

## Materials and Methods

### Study site and sampling procedure

The study area located at the eastern Qinghai-Tibetan Plateau with the average altitude above 3000 m a.s.l. Specifically, the study site was located at Oula village of the Maqu Wetland Protection Area (E 100°45′∼102°29′, N 33°06′∼34°30′) in Gansu Province, China. The mean daily air temperature is 1.2°C, with the lowest mean air temperature, -10°C, in January and highest mean air temperature, 11.7°C, in July. Mean annual precipitation is 620 mm and mainly occur during the summer. The grazing pastures for the animals is typical alpine meadow,the main vegetative cover is as follows: *Kobresia kansuensis,Thalictrum aquilegifolium var. sihiricum,Stipa capillata, Potentilla fragarioides, Saussurea hieracioides, Taraxacum mongolicum, Anemone baicalensis var. kansuensis, Anemone rivularis var. flore-minore, Euphorbia esula, Medicago ruthenica, Plantago asiatica*. Yak and Tibetan sheep, even with some wild animals, are living together and grazing at the same pastures without any additional feeding. The yak population is around 200 with the ages ranging between 1–3 years old. There are 300 Tibetan sheep with the ages between 1– 1.5 years old.

Sampling procedures were performed twice, on 29th March,2016 as for spring season samples, after the melting of snow and before the sprouting of grass, and on 3rd November, 2016 for autumn season samples, approximately the fattest time of the animals. Fresh fecal samples were collected in the early morning. Samples were put into the sterilized plastic tubes and kept in the liquid nitrogen until further experimental analyses. In total, 226 fresh fecal samples were collected, including 136 yak fecal samples – 56 in spring (thereafter coded as SprY) and 80 in autumn (AutY), and 90 Tibetan sheep fecal samples - 43 in spring (SprS) and 47 in autumn (AutS).

All of the experimental protocols and procedures were approved by the Institutional Animal Care and Use Committee of Lanzhou Institute of Husbandry and Pharmaceutical Science of Chinese Academy of Agricultural Sciences (Approval No. NKMYD201611). Animal welfare and experimental procedures were performed strictly in accordance with the Guidelines for the Care and Use of Laboratory Animals issued by the US National Institutes of Health.

### DNA extraction, PCR amplification, and high-throughput sequencing

Genomic DNA was extracted from the fecal samples by using TIANGEN DNA Stool Mini Kit (TIANGEN, cat#DP328) following the manufacturer’s instructions. The quality and quantity of extracted DNA were assessed by NanoDrop ND-1000 spectrophotometer (NanoDrop Technologies, Wilmington, USA). The V3-V4 region of the 16S rRNA gene was amplified using barcoded primers. Amplicons were extracted from 2% agarose gels and purified using the AxyPrep DNA Gel Extraction Kit (Axygen Biosciences, Union City, CA, U.S.) according to the manufacturer’s instructions and quantified using QuantiFluor(tm) -ST (Promega, U.S.). Purified amplicons were pooled in equimolar and paired-end sequenced (2 × 300 bp) on an Illumina MiSeq platform according to the standard protocols. The sequencing procedures were delegated to the commercial company, Gansu GeneBioYea Biotechnology Co. Ltd.

### Sequencing data processing

Raw FASTQ files were de-multiplexed and quality-filtered using QIIME (version 1.9.1) [19] with the following criteria: (i) The 300-bp reads were truncated at any site that obtained an average quality score of < 20 over a 10-bp sliding window, and the truncated reads shorter than 50 bp were discarded; (ii) exact barcode matching, two nucleotide mismatch in primer matching, and reads containing ambiguous characters were removed; (iii) only overlapping sequences longer than 10 bp were assembled according to their overlapped sequence. Reads that could not be assembled were discarded. Operational taxonomic units (OTUs) with 97% similarity cutoff were clustered using UPARSE (version 7.1)[20], and chimeric sequences were removed using UCHIME [21]. The taxonomy of each 16S rRNA gene sequence was analyzed against the greengene database at a confidence threshold of 70%, respectively. The rarefaction analysis based on Mothur v.1.35.1 (https://www.mothur.org) was conducted to reveal the diversity indices, including the ACE, Chao1, Shannon, Simpson, and coverage indices [22].

## Data analyses

Two-way ANOVA was utilized to explore the effects of the season and host species on richness, evenness, and diversity of microbial communities. Beta diversity of Bray-cutis distance between the samples in the same groups and between different grouped were analyzed and box-plot was generated to show the differences. Principal coordinates analysis (PCoA) plots were generated to compare the composition of bacterial/archaeal community structure among different treatments. Permutational multivariate analysis of variance (PERMANOVA) on the Bray-Curtis metric produced by PCoA analysis was performed to test the significant difference in community composition among the treatments. All the above analyses were completed by R (versions 3.3.3, R Core Team. 2016). Non-parametric ANOVA analysis was conducted with ‘lmPerm’ package (Wheeler and Torchiano 2016), multivariate analyses were conducted with ‘vegan’ package (Oksanen et al. 2016). Phylogenetic Investigation of Communities by Reconstruction of Unobserved States (PICRUSt) was utilized to predict metagenome functional content from the 16S rRNA gene surveys (http://picrust.github.com) [23]. Venn diagrams were constructed to show the unique or shared OTUs and also the KEGG functional genes predicted by PICRUSt.

## Acknowledgements

The work was supported by the earmarked fund for the China Agriculture Research System (No.CARS-38) and the Central Public-interest Scientific Institution Basal Research Fund (No.1610322017014).

